# An optimized electroporation approach for efficient CRISPR/Cas9 genome editing in murine zygotes

**DOI:** 10.1101/281402

**Authors:** Simon E. Tröder, Lena K. Ebert, Linus Butt, Sonja Assenmacher, Bernhard Schermer, Branko Zevnik

## Abstract

Electroporation of zygotes represents a rapid alternative to the elaborate pronuclear injection procedure for CRISPR/Cas9-mediated genome editing in mice. However, current protocols for electroporation either require the investment in specialized electroporators or corrosive pre-treatment of zygotes which compromises embryo viability. Here, we describe an easily adaptable approach for the introduction of specific mutations in C57BL/6N mice by electroporation of intact zygotes using a common electroporator with synthetic CRISPR/Cas9 components and minimal technical requirement. Direct comparison to conventional pronuclear injection demonstrates significantly reduced physical damage and thus improved embryo development with successful genome editing in up to 100% of living offspring. Hence, our novel approach for <u>E</u>asy <u>E</u>lectroporation of <u>Zy</u>gotes (EEZy) allows highly efficient generation of CRISPR/Cas9 transgenic mice while reducing the numbers of animals required.

## Introduction

To understand the mechanisms underlying human diseases biomedical research relies on the availability of animal models with precise recapitulation of human genetics. To this end, state-of-the-art genome editing by use of the CRISPR/Cas system (clustered regularly interspaced short palindromic repeats/CRISPR-associated protein) in mouse zygotes has enabled the fast and scarless introduction of specific mutations with high efficiency [1-4]. The most commonly used CRISPR/Cas system consists of three basic components: the endonuclease Cas9, the sequence specific crRNA (CRISPR RNA) and the generic tracrRNA (trans-activating crRNA). Naturally occurring as a paired guide RNA (pgRNA) the two sequences have been artificially fused to a single guide RNA (sgRNA) molecule for practical reasons [3]. At the desired locus, CRISPR/Cas9-mediated DNA double strand breaks are either repaired by non-homologous end joining (NHEJ), resulting in random insertions and deletions (INDELs) or, if a DNA repair template is provided, by homology directed repair (HDR). The latter mechanism can be exploited to introduce site-specific mutations into the locus of interest. While CRISPR/Cas9-mediated transgenesis has emerged as the novel standard technology for the generation of mouse models, the delivery of the necessary components into zygotes still relies on technically demanding and invasive pronuclear injection (PNI). Recently, three independent groups demonstrated electroporation of zygotes as an alternative delivery route for Cas9 mRNA, guide RNAs and short single-stranded oligonucleotides (ssODN) as DNA repair templates for precise CRISPR/Cas9-mediated transgenesis [5-7]. Subsequently, electroporation of pre-assembled Cas9/sgRNA ribonucleoproteins (RNP) proved to drastically enhance transgenic efficiencies [8, 9]. However, several obstacles within the current protocols hinder the broader application of electroporation: Firstly, they either require the costly investment in a specialized electroporator device, or use corrosive pre-treatment of the embryo by acidic Tyrode’s solution to allow uptake of the CRISPR/Cas9 components. Weakening of the zona pellucida, a vital part of the embryo, by acidic Tyrode’s solution compromises embryo viability [10-12] while the necessary incubation time of as short as 10 s [7] is challenging to be reproducibly applied even for highly experienced users. Furthermore, the batch specific activity of acidic Tyrode’s solution [8] challenges its safe and reproducible application. Secondly, synthetic guide RNAs have recently become commercially available circumventing laborious preparation of *in vitro* transcribed sgRNA. However, the impact of delivering these synthetic guide RNAs by electroporation on embryo viability has not been assessed side-by-side with conventional sgRNA. Thirdly, major results optimizing the electroporation conditions are based on experiments with zygotes from robust F1 hybrid mice whereas the vast majority of research questions demand transgenesis in the more sensitive but most commonly used inbred C57BL/6N mice in order to keep the results comparable and reproducible [6, 7, 13-15].

To overcome these constraints, we combined the current data on electroporation in order to develop an easily adaptable protocol for efficient CRISPR/Cas9-mediated transgenesis of C57BL/6N zygotes with minimal technical demand. We aimed to omit any treatment weakening the zona pellucida and to establish the protocol on a standard electroporation device. Utilization of commercially available CRISPR/Cas9 components, most importantly synthetic guide RNAs, further simplified the process and enhanced the reproducibility as well as the embryo viability. We found that our improved CRISPR electroporation, which we refer to as EEZy (<u>E</u>asy <u>E</u>lectroporation of <u>Zy</u>gotes), enables the introduction of specific mutations as efficient as delivery by conventional PNI while it significantly enhances embryo development *in vitro* and live birth rates *in vivo*. Genome editing accomplished in up to 100% of the founder mice demonstrate complete penetration of zygotes without any weakening of the zona pellucida. Thus, our EEZy approach represents an easily adaptable and broadly applicable technique for CRISPR-mediated mouse transgenesis which outperforms conventional PNI.

## Materials and methods

### Ethics statement

All mouse protocols were in accordance with European, national and institutional guidelines and approved by the State Office of North Rhine-Westphalia, Department of Nature, Environment and Consumerism (LANUV NRW, Germany; animal study protocol AZ 84-02_04_2014_A372). Mice were kept in the specific and opportunistic pathogen free animal facility of the CECAD Research Center of the University of Cologne at 22°C (± 2°C) and a humidity of 55% (± 5%) under 12 h light cycle with access to water and food *ad libitum*. Mice were anesthetized with ketamine (Ketaset, Zoetis) and xylazine (Rompun, Bayer) and euthanized by cervical dislocation. Carprofen (Rimadyl, Zoetis) was used as analgesic after surgery. All efforts were made to minimize suffering.

### Generation of guide RNAs

Custom crRNA (IDT, Alt-R™ crRNA) and generic tracrRNA (IDT, 1072532) were resuspended to 100 µM in sterile and nuclease-free T_10_E_0.1_ buffer (10 mM Tris-HCl, 0.1 mM EDTA, embryo-tested water (Sigma, W1503)) prepared as described [16]. 50 µM crRNA:tracrRNA complexes (pgRNA) were generated by subjecting equimolar ratios to 95°C for 5 min followed by reduction of 5°C/min. sgRNA was generated by T7 RNA polymerase mediated *in vitro* transcription (NEB, E2040S) from the pSpCas9(BB)-2A-GFP (PX458) plasmid (Addgene #48138) from Feng Zhang and column purified (Qiagen, 217004). Guide RNAs were stored at −80°C. crRNA sequences are listed in S1 Table.

### Mouse transgenesis

A step-by-step protocol for EEZy is available on protocols.io (dx.doi.org/10.17504/protocols.io.ndzda76). Unless otherwise stated, mouse pre-implantation embryos were kept in pre-incubated M16 in a CO_2_ incubator (5% CO_2_, 37°C, 95% humidity; Labotect C16) or handled in M2 outside the CO_2_ incubator. M16 and M2 were prepared as described [17]. C57BL/6NRj zygotes for electroporation were either directly purchased from Janvier Labs (SE-ZYG-CNP) or collected from C57BL/6NRj females. Therefore, 3-4-week-old donor females (i.e. 12-14 g body weight) were superovulated 72 h prior zygote collection by intraperitoneal administration of 5 IU of pregnant mare serum gonadotrophin (ProSpec, HOR-272 or Aviva Systems Biology, OPPA01037) followed by 5 IU of human chorionic gonadotrophin (Intervet, Ovogest) 48 h later. 0.5 days post coitum (dpc) zygotes were collected from the oviducts of donor females upon 1:1 mating with C57BL/6NRj males as described in published protocols [17]. If indicated C57BL/6NRj zygotes were collected upon *in vitro* fertilization without using reduced glutathione essentially as described in published protocols with the minor modification of using Cook RVFE (Cook Medical, K-RVFE-50) as fertilization medium and M16 for post fertilization wash steps instead of human tubal fluid media [17]. For electroporation of zygotes Cas9 RNPs were assembled by combining 4 µM Cas9 protein (IDT 1074181) with 4 µM of assembled pgRNA and sgRNA, respectively, in 20 µl Opti-MEM (Thermo Fisher Scientific, 31985062) and incubation for 10 min at room temperature. If applicable, 10 µM ssODN (IDT, custom Ultramer Oligo; sequences are listed in S1 Table) or 200 ng/µl targeting vector were added. The plasmid targeting vector for integration of a splice-acceptor fused Venus reporter in the *Gt(ROSA)26Sor* locus was a kind gift from Ralf Kühn [16]. If indicated, the Cas9 RNPs were generated as a 2× solution (8 µM Cas9 protein; 8 µM gRNA) in freshly prepared sterile 2× pre-mix (20 mM HEPES pH7.5 (Sigma, H3375), 150 mM KCl (Sigma, P9333), 1 mM MgCl2 (Sigma, 63068), 10% glycerol (Sigma, 49767) and 1 mM reducing agent TCEP (tris(2-carboxyethyl)phosphine (Sigma, C4706), nuclease-free H_2_O (Qiagen, 129114)), incubated for 10 min at room temperature and add to 20 µM ssODN in 10 µl Opti-MEM. Each Cas9 RNP mix was kept on ice until use. For electroporation, zygotes were washed in batch through 5 drops of M2. After wash through one drop of Opti-MEM the zygotes were transferred with as little media as possible to 20 µl of Cas9 RNP mix. Using a P20 pipette, the entire solution was transferred to a pre-warmed 1 mm cuvette (BioRad, 1652089) and placed in the BioRad Gene Pulser XCell electroporator. Two square wave pulses were applied (30V, 3 ms pulse duration, 2 pulses, 100 ms interval). The zygotes were retrieved from the cuvette using a P100 pipette and two flushes of 100 µl M16 and transferred to a culture dish (Falcon, 353037) containing 500 µl M16. If indicated zygotes were treated with acidic Tyrode’s solution (Sigma, T1788) for 10 s and the reaction stopped by adding M2. Zygotes for PNI were exclusively collected from C57BL/6NRj females upon natural mating as described above. After visual inspection for the presence of two pronuclei zygotes were placed in a paraffin oil covered M2 containing injection chamber. Microinjection was performed using an Axio Observer.D1 microscope (Zeiss) and microinjector devices CellTram and FemtoJet with TransferMan NK2 micromanipulators (Eppendorf). The CRISPR/Cas9 solution was injected into the male pronucleus using injection capillaries (BioMedical Instruments, BM100F-10; type PI-1.6). If not stated otherwise the CRISPR/Cas9 injection mix was prepared as described for electroporation but containing 400 nM gRNA, 200nM Cas9 protein, 30 ng/µl Cas9 mRNA (TriLink, L-6125-20) and 500 nM ssODN. Microinjected zygotes were transferred into 500 µl M16 and lysed embryos were removed approximately 1 h after injection. On the next day, 2-cell stage embryos were transferred unilateral into oviducts of pseudo-pregnant 0.5 dpc RjHan:NMRI females or continued to culture until the blastocyst stage at 3.5 dpc.

### Genotyping analysis

DNA of individual blastocyst stage embryos was extracted in 10 µl of QuickExtract DNA extraction solution (Epicentre, QE09050) in a thermocycler at 65°C for 15 min followed by 95°C for 15 min and stored at −80°C until use. Except depicted otherwise, two sequential rounds of PCR with Herculase II fusion DNA polymerase (Agilent Technologies, 600677) were performed according to the manufacturer’s instruction using 2 µl DNA template. The primer sequences and thermocycling conditions are listed in S1 Table. For restriction fragment length polymorphism (RFLP) assay the PCR amplicons were column purified (Macherey-Nagel, 740609.250) or used right away for digestion with the indicated restriction enzyme (NEB) and the DNA fragments were analyzed by agarose gel electrophoresis. Percentage value of embryos harboring the desired mutation were counted by the presence of digested DNA fragments. Ear biopsies were lysed in 75 µl alkaline lysis buffer (25 mM NaOH, 0.2 mM EDTA) for 30 min at 95°C followed by 75 µl neutralization solution (40mM Tris-HCl). Sanger sequencing of PCR amplicons was performed by the Cologne Center for Genomics (CCG).

### Statistical analysis

For calculation of statistical significance, standard deviation and arithmetic mean of differences Prism (GraphPad) and Excel (Microsoft) were employed. Box plots were generated by Prism. Statistical significance was assessed using a two-tailed, unpaired Student’s T-test. Differences were considered significant below a *p*-value of 0.05.

## Results

In order to establish an easily adaptable protocol for CRISPR/Cas9 electroporation in C57BL/6N zygotes we intentionally utilized a standard electroporator (BioRad Gene Pulser XCell). Similar devices are broadly available in biomedical laboratories. Two pulses of 3 ms at 30 V were previously demonstrated as the ideal balance between editing efficiency and embryo survival for electroporation with this device [8]. We chose to electroporate 4 µM of pre-assembled Cas9 RNPs as this concentration does not impact embryo viability and Cas9 protein has been shown to dramatically outperform Cas9 mRNA electroporation regarding transgenic efficiency [8, 9].

To validate our EEZy approach, we introduced a new BsaI restriction site in the *Nphs2* gene by electroporation of solely commercially available CRISPR/Cas9 components (guide RNA, Cas9 protein and ssODN repair templates) and analyzed the genotype of the blastocysts by RFLP analysis. In a first set of experiments, we determined whether previously suggested additives to the electroporation buffer are necessary to maintain editing efficiency and embryo development [8]. We compared the ratio and genotype of developed blastocysts upon EEZy in conventional electroporation buffer containing freshly added additives (Opti-MEM supplemented with reducing agent TCEP, HEPES, KCl, MgCl2 and glycerol) with blastocysts from zygotes electroporated only with commercially available Opti-MEM as electroporation buffer. Three independent experiments showed no advantage of electroporation buffer additives for transgenesis, analyzed by RFLP analysis (Fig 1A and 1B) and confirmed by Sanger sequencing (S1 Fig), and embryonic development (Fig 1C). The consistently high HDR-efficiency in these experiments also validated that pre-treatment of zygotes with Tyrode’s solution is indeed dispensable for efficient transgenesis in our EEZy approach. In confirmation, we detected equally high HDR-efficiencies when we targeted another genetic locus (*Atp1a1*) using our EEZy approach. Pre-treatment of zygotes with acidic Tyrode’s solution in this experiment did not enhance insert integration (S2 Fig).

**Fig 1.**
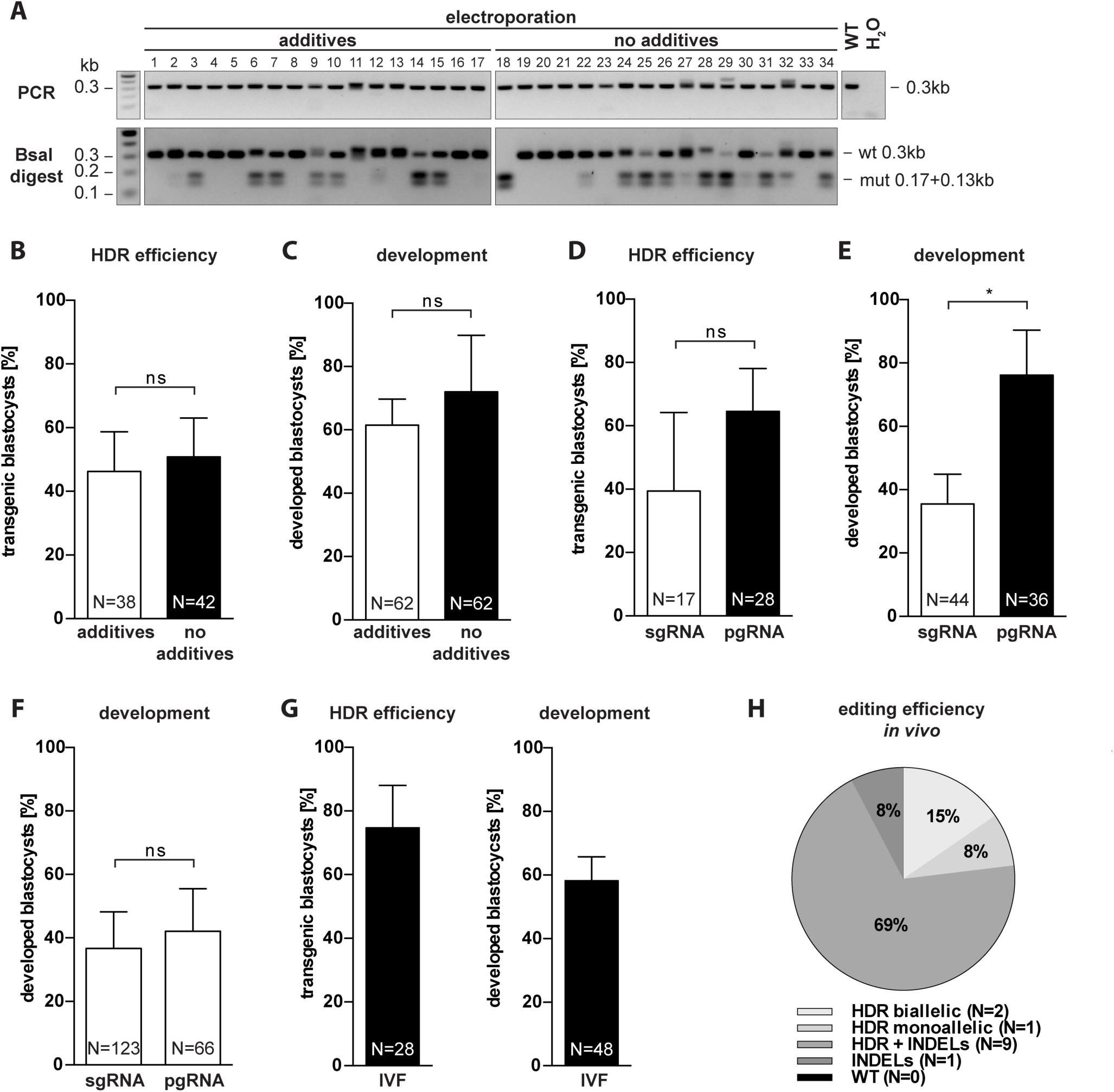
Characterization of EEZy for specific genome editing in intact C57Bl/6N zygotes. (A) Representative RFLP analysis for evaluation of the HDR efficiency in *Nphs2*-targeted blastocysts upon EEZy using Opti-MEM with and without additives and (B) quantification of the genotypes from three independent experiments. PCR controls from untreated blastocysts (WT) and without DNA template (H_2_O) are depicted. (C) Assessment of embryo development from (B) as percentage of developed blastocysts from zygotes after EEZy. (D) Quantification of RFLP analysis from *Nphs2*-targeted blastocysts upon EEZy using sgRNA or pgRNA from three independent experiments and (E) assessment of percentage of developed blastocysts from these zygotes. (F) Assessment of developed blastocysts after PNI using pgRNA or sgRNA targeting *Nphs2* from four independent experiments. (G) Quantification of RFLP genotyping at the *Nphs2* locus and percentage of developed blastocysts after EEZy of zygotes obtained by IVF from three independent experiments. (H) Quantification of Sanger sequencing of biopsies from *Tmem218* transgenic mice generated by EEZy. Data of a total of 13 mice displaying either solely the desired mutation (HDR), a mixture of the desired mutation and INDELs or only INDELs are depicted. Data are means ± standard deviation. *p < 0.05, ns = non-significant. N = total number of embryos analyzed.

To circumvent laborious *in vitro* transcription of sgRNA we next utilized synthetic guide RNA consisting of paired crRNA:tracrRNA complexes (pgRNA). The use of these commercially available pgRNAs for electroporation has recently been described but their impact on editing efficiency and embryo development compared to conventional sgRNA is unknown [18]. We did not detect a significant difference in HDR efficiency between EEZy with either sgRNA or pgRNA targeting *Nphs2* (Fig 1D). EEZy with sgRNA, however, clearly decreased embryo viability in our experiments (Fig 1E). This negative impact on embryo development was not observed when we compared the use of these types of guide RNAs in PNI (Fig 1F).

We also assessed whether our EEZy approach is effective for delivery of plasmid targeting vectors to generate CRISPR-mediated large genomic integrations as such has not been published to date using electroporation. We attempted to integrate a Venus reporter transgene by electroporation of Cas9 RNPs targeting the *Gt(ROSA)26Sor* locus and a plasmid vector already successfully used by conventional PNI [16]. Despite several efforts, we could not generate blastocysts with integration of the transgene with our approach even though INDELs were detected at high frequencies (S3 Fig).

Zygotes for electroporation can either be collected from donor females upon natural mating or generated by IVF from oocytes. The potential of the latter zygotes to reduce mosaicism in targeted mice due to the feasibility of transgenesis in early pronuclear zygotes has recently been demonstrated [13]. Using our EEZy approach with IVF generated zygotes we were able to obtain transgenic embryos with both, high editing efficiency and developmental rates (Fig 1G), comparable with zygotes from natural mating. Notably, we did not weaken the zona pellucida of IVF zygotes with reduced glutathione which has previously been suggested to allow delivery of CRISPR components by electroporation [9]. Taken together, our approach is independent of the zygote source and proves that zygotes with intact zona pellucida can be edited with high efficiency in a standard electroporator. Consistently, we successfully generated living mice with integration of a point mutation in the coding region of another gene (*Tmem218*) employing our EEZy approach (Fig 1H). Sequencing revealed HDR editing in up to 92% of the founder mice, with 23% of animals harboring solely the desired mutation (15% biallelic, 8% monoallelic), validating our EEZy approach for generation of specific mutant mice *in vivo*. Strikingly, genome editing occurred in 100% of the offspring which proves, once again, that corrosive zona weakening is dispensable during EEZy.

Next, we aimed to test whether our EEZy approach would increase the number of embryos amenable to genome editing. Conventional PNI relies on the presence of visible pronuclei. In our experience, this fraction represents about 70% of all harvested zygotes (Fig 2A). The competency of zygotes without visible pronuclei for genome editing has not yet been demonstrated. We electroporated zygotes without visible pronuclei and compared the editing efficiency at the *Nphs2* gene to the results of zygotes without pre-selection of pronuclei. Indeed, the developed blastocysts of either zygote fraction showed similar rates of HDR editing (Fig 2B and 2C). However, significantly fewer zygotes without visible pronuclei developed into morphologically intact blastocysts (Fig 2D). Hence, electroporation enables, in principal, transgenesis of zygotes non-accessible to conventional PNI although embryo development seems impaired in this fraction of zygotes.

**Fig 2.**
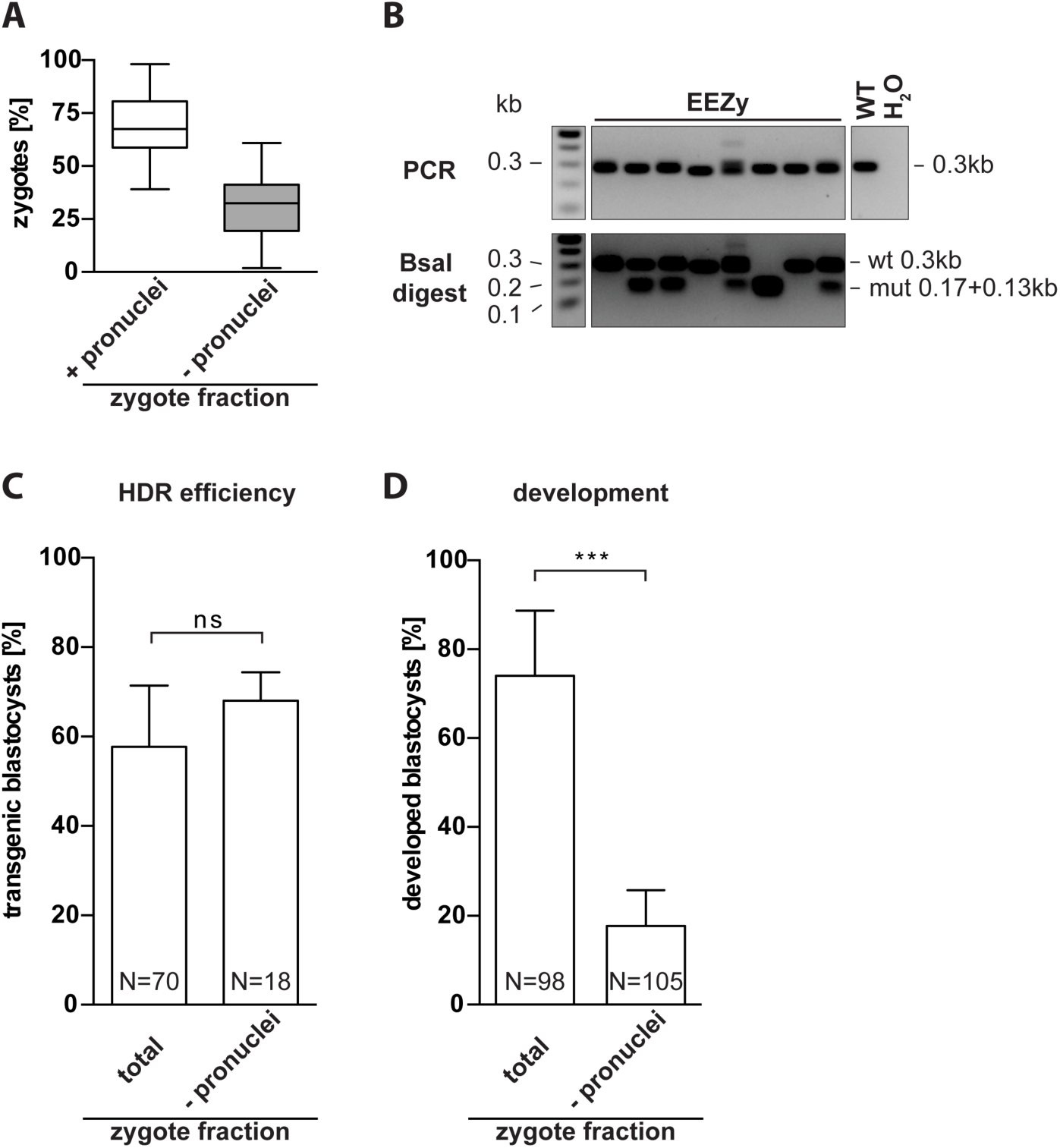
Electroporation of zygotes incompatible with pronuclear injection. (A) Average ratio of harvested zygotes from natural mating with visible pronuclei (+) and without visible pronuclei (-) from 47 independent experiments. Total number of embryos analyzed was 6368 and 3314 for zygotes with and without visible pronuclei, respectively. (B) Representative RFLP analysis for evaluation of HDR efficiency in *Nphs2*-targeted blastocysts upon EEZy of zygotes without visible pronuclei. PCR controls from untreated blastocysts (WT) and without DNA template (H_2_O) are depicted. (C) Quantification of the genotype from three independent experiments using zygotes without visible pronuclei are compared to EEZy of all zygotes from the experiments depicted in Fig 1B and 1D, right columns. (D) Percentage of developed blastocysts from zygotes after EEZy from (C). Box plot represents median, boxes equal 25 to 75 percentiles, whiskers include all values. Remaining data are means ± standard deviation. ***p < 0.001, ns = non-significant. N = total number of embryos analyzed.

Previous studies comparing the impact on embryo viability of CRISPR component delivery by electroporation and PNI gained inconsistent results. While most studies clearly showed that electroporation outperformed conventional PNI regarding embryo development or live birth rate [5, 7, 8, 14] there was also some evidence for mixed outcomes [19, 20]. Despite this tendency no statistical hypothesis testing has been possible due to changing parameters within the experiments of the individual studies. To systematically assess the effect of our EEZy approach on embryo development we compared untreated with electroporated zygotes regarding their capacity to form blastocysts *in vitro* (Fig 3A). Neither EEZy *per se* (Mock) nor EEZy of CRISPR components significantly affected embryo development. Next, we directly compared the results of our EEZy approach to transgenesis by conventional PNI using the same knock-in strategy for *Nphs2* as before. In line with previously published results, on average 82% of the zygotes survive in our PNI (Fig 3B) [21] while about 95% of zygotes were recovered upon EEZy. We have rarely seen immediate lysis of zygotes after EEZy which is a known phenomenon upon PNI and responsible for the lost zygotes during our PNI (Fig 3B). In contrast, the minor fraction of zygotes lost during EEZy only account for embryos which typically cannot be retrieved from the electroporation cuvette. Consistent with the assumption that EEZy is less damaging we observed more of the recovered zygotes from EEZy to develop to blastocysts as compared to PNI (Fig 3C). The combined results of reduced embryo toxicity (Fig 3B) and higher developmental capacity (Fig 3C) summarized in Fig 3D clearly emphasize the potential of EEZy to significantly lower the numbers of zygote donors. Like published before delivery by electroporation does not compromise the editing efficiency of the Cas9 RNP in comparison to PNI (Fig 3E and S4 Fig) [14, 19]. To confirm the capability of EEZy to reduce animal numbers *in vivo* we compared the number of offspring upon transgenesis at different genetic loci to the results from conventional PNI. Consistent with our findings of improved pre-implantation development we uniformly gained higher numbers of offspring with EEZy as compared to PNI (Table 1).

**Table 1.**
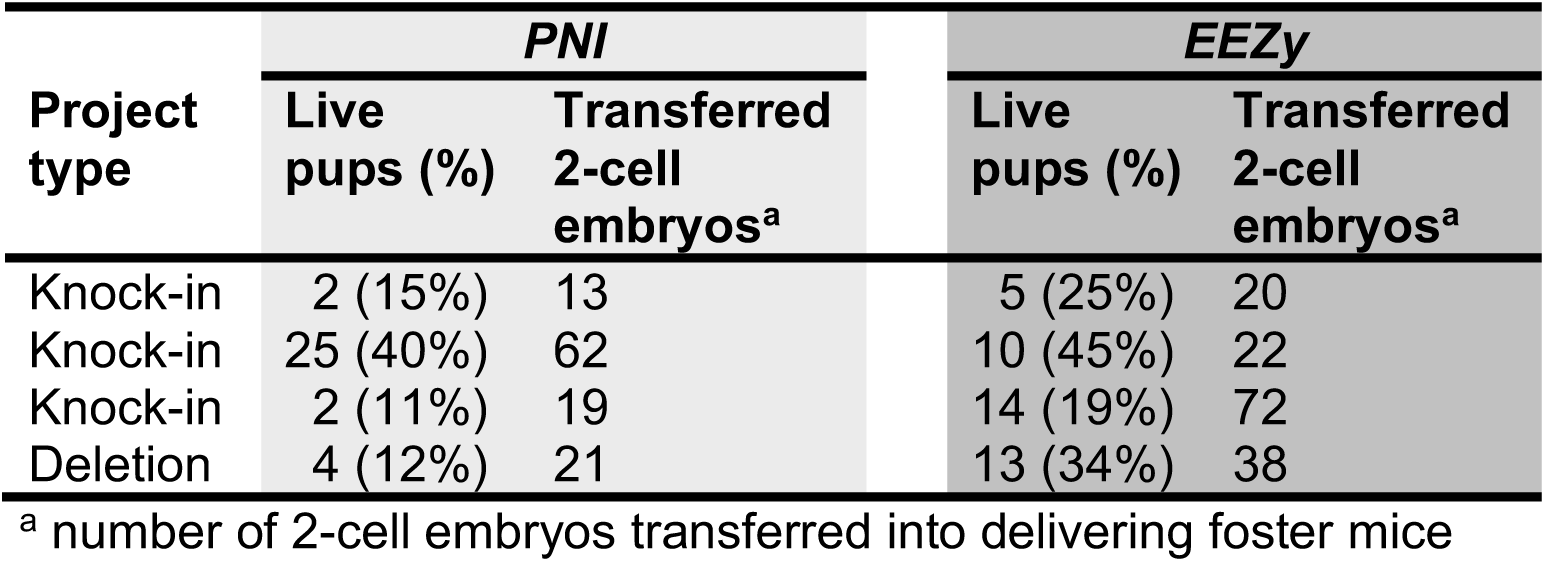
Live birth rate of offspring upon transgenesis by PNI or EEZy.

| Project type | PNI |  | EEZy |  |
| --- | --- | --- | --- | --- |
|  | Live pups (%) | Transferred 2-cell embryos<sup>a</sup> | Live pups (%) | Transferred 2-cell embryos<sup>a</sup> |
| Knock-in | 2 (15%) | 13 | 5 (25%) | 20 |
| Knock-in | 25 (40%) | 62 | 10 (45%) | 22 |
| Knock-in | 2 (11%) | 19 | 14 (19%) | 72 |
| Deletion | 4 (12%) | 21 | 13 (34%) | 38 |

<sup>a</sup> number of 2-cell embryos transferred into delivering foster mice

**Fig 3.**
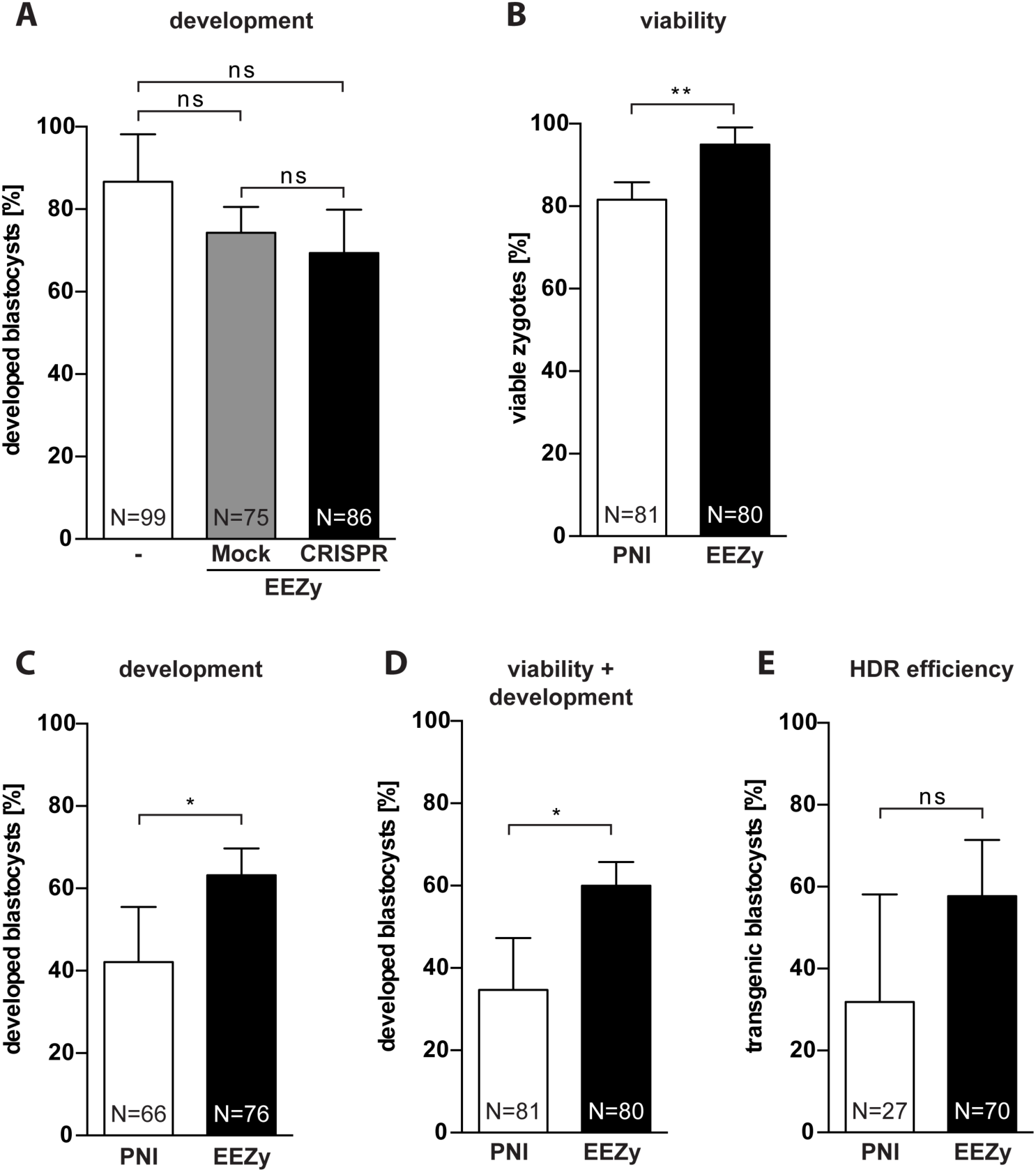
Embryo toxicity of EEZy as compared to pronuclear injection. (A) Assessment of embryo development as percentage of developed blastocysts from zygotes after EEZy. Untreated zygotes are compared to zygotes electroporated with Opti-MEM (Mock) or CRISPR/Cas9 components targeting the *Gt(ROSA)26Sor* locus (CRISPR). Data represent three independent experiments. (B) Quantification of viable *Nphs2*-targeted zygotes from four experiments immediately after PNI compared to electroporated zygotes and the correspondingly developed blastocysts (C). (D) Developed blastocyst calculated from the number of zygotes before transgenesis in (B). Quantification of RFLP genotyping from four independent experiments for HDR efficiency in blastocysts from zygotes after PNI compared to electroporated zygotes from the experiments depicted in Fig 1B and 1D, right columns. Data are means ± standard deviation. *p < 0.05, **p < 0.01, ns = non-significant. N = total number of embryos analyzed.

Taken together, our novel EEZy approach enables efficient introduction of specific mutations with minimal technical demand and high embryo viability.

## Discussion

In this study, we established a novel and facile method to use electroporation for the generation of CRISPR/Cas9-based transgenic mouse lines. Basically, this approach uses synthetic CRISPR components in a standard electroporation device with intact zygotes either from natural mating or from IVF. We demonstrate that zona pellucida weakening is dispensable for efficient generation of mutant mice *in vivo* by our EEZy approach. Pre-treatment of zygotes by acidic Tyrode’s solution has been initially established to increase the permeability of the zona pellucida for large RNA molecules during electroporation and has been subsequently shown to be necessary for gene editing using Cas9 mRNA of about 4,500 nucleotides length [7, 22]. Uptake of small molecules like morpholinos of only 25 nucleotides length, however, neither required nor greatly benefited from acidic Tyrode’s pre-treatment of zygote prior to electroporation [12]. We accomplished highly efficient gene editing in embryos and living offspring by EEZy with Cas9 RNP complexes and small DNA repair templates of about 150 nucleotides length. The sufficient permeability of intact zygotes in our experiments may be explained by the compact nature of the Cas9 RNP complex in contrast to large Cas9 mRNA. Recently, long ssODN repair templates of about 1,500 nucleotides length have been successfully used for generation of mutant mice by CRISPR-PNI [23]. Assuming low permeability of intact zygotes for large nucleic acids, it is conceivable that zona pellucida weakening might still be valuable for electroporation of these long ssODNs. For the same purpose, the pulse frequency or duration during EEZy may be increased to facilitate enhanced uptake of nucleic acids. Although the current parameters have been identified as the ideal balance between editing efficiency and embryo survival previously [8] it has been demonstrated that gene editing of difficult targets may benefit from increased pulse frequencies or durations. However, in this case lower embryo viability is inevitable [6, 9, 20].

Our results revealed significant less embryo toxicity of synthetic pgRNAs as compared to sgRNAs from *in vitro* transcription during our EEZy approach. In confirmation with previous studies we did not detect this difference using conventional PNI [24]. We hypothesize that the ten times higher amounts of guide RNA needed during EEZy may account for the elevated embryo toxicity of sgRNAs during EEZy. Although it remains interesting whether the nature of the sgRNA or traces from its generation are responsible for the embryo toxicity our data validate synthetic pgRNAs as less harmful during electroporation. While our manuscript was in preparation another group independently used synthetic pgRNAs for successful generation of CRISPR/Cas9-mediated mouse mutants via electroporation [18]. Although they did not compare pgRNA and sgRNA side-by-side their data nicely confirm our finding that synthetic pgRNA is indeed an efficient and safe alternative to conventional sgRNA.

Successful CRISPR-mediated generation of large genomic integrations by electroporation of plasmid targeting vectors into mouse zygotes has not been published to date. Interestingly, an equivalent attempt of such integration by electroporation most recently also failed in rat zygotes [25]. The data suggested that plasmid vectors do not reach the nucleus although they were detected in the cytosol of the embryo upon electroporation. We therefore speculate that insufficient permeability of the nuclear membrane may also be responsible for the fact that we did not detect any transgene integration upon EEZy with a plasmid targeting vector.

The presence of visible pronuclei, a requirement for conventional PNI, is dispensable for EEZy. Our data demonstrate for the first time that blastocysts from zygotes without visible pronuclei display the same HDR editing rates as blastocysts from zygotes with visible pronuclei. When performing CRISPR-PNI in our daily routine we therefore electroporated this PNI-incompatible zygote fraction to evaluate the guide RNA efficiency for future projects in *in vitro* blastocyst assays. In these assays targeted zygotes are cultured to the blastocyst stage and genotyped by PCR [26].

We further found a significantly lower toxicity of our EEZy approach in direct comparison to PNI. While delivery of CRISPR components by PNI has been reported to induce physical damage, and impede embryo development [21], EEZy had virtually no impact on embryo development and subsequently resulted in higher numbers of offspring. These findings are in line with published results of improved pre-implantation development from B6D2F1 hybrid zygotes [14] and higher live birth rates of inbred C57BL/6J mice from zygotes after CRISPR electroporation as compared to PNI [8]. In addition, we show for the first time that higher numbers of embryos can be retrieved after electroporation due to less acute damage observed during PNI. In fact, we have hardly ever observed immediate lysis of embryos employing EEZy which may additionally result from the higher resistance of intact zygotes compared to zona pellucida weakened zygotes [12]. Along this line, our data demonstrate the potential of EEZy to profoundly decrease animal numbers needed for the generation of transgenic mice. Reduction of animal numbers is one of the key aspects of the 3R principles (Replacement, Reduction, Refinement) for good animal welfare in research [27]. During CRISPR-mediated transgenesis, either by electroporation or PNI, animals are needed for collecting zygotes and as recipient foster females for the targeted embryos. Due to enhanced embryo viability and development EEZy lowers the demand for both, the zygote donors and recipient females in comparison to conventional PNI highlighting the value of this technique for more humane animal research.

In summary, with EEZy we established a simplified and easily adaptable electroporation procedure using intact C57BL/6N zygotes for highly efficient CRISPR/Cas9-mediated generation of specific mutant mice with minimal impact on embryo viability.

## Acknowledgement

We thank Patrick Jankowski, Kerstin Weisheit and Stefanie Wasserburger for excellent technical support. We thank Ralf Kühn for providing us with expression vectors and Anna Johann for critical reading of the manuscript.

