## Supplementary Materials for "An optimized electroporation approach for efficient CRISPR/Cas9 genome editing in murine zygotes"

### Supporting information

**S1 Figure**

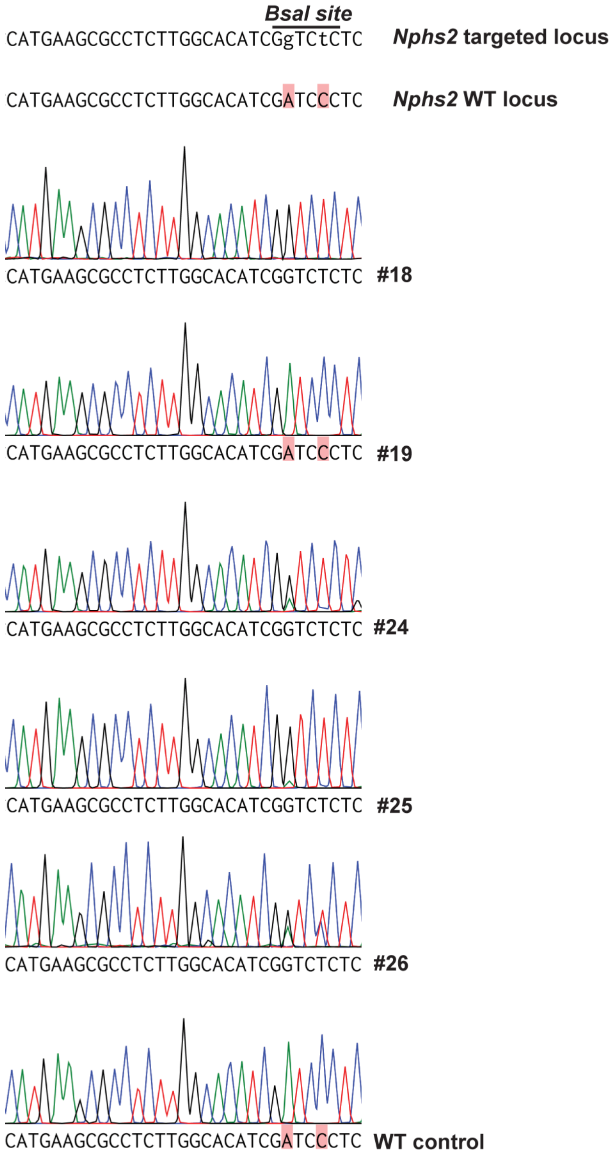


**S1 Fig. Sanger sequencing of representative *Nphs2*-targeted blastocysts.** Sequencing results of the indicated blastocysts from Fig 1A. The expected DNA sequence of either the endogenous WT locus or the desired mutation (new BsaI restriction site) is depicted at the top.

**S2 Figure**

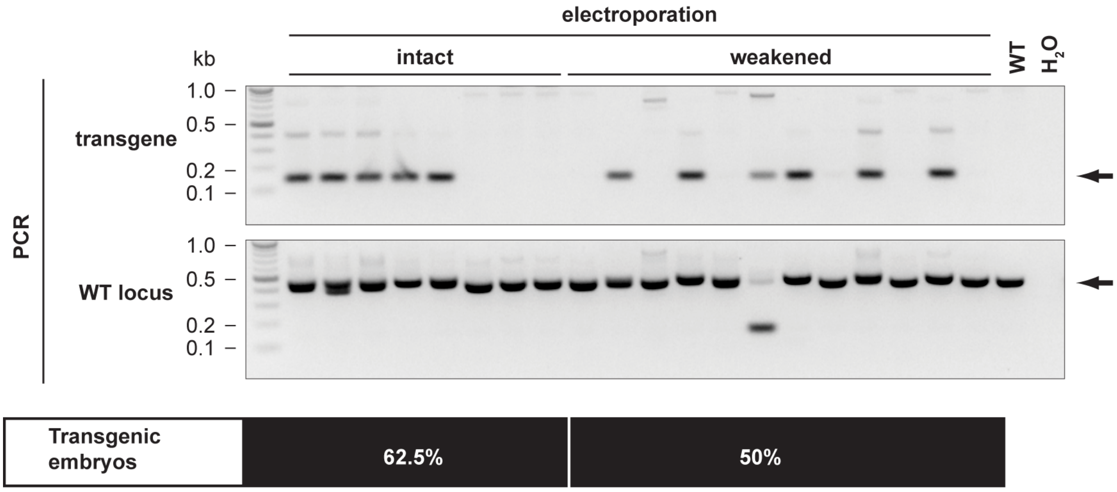


**S2 Fig. Acidic Tyrode's treatment of zygotes prior electroporation.** Myc tag insertion into the *Atp1a1* gene by electroporation. PCR genotyping and the percentage of transgenic 3.5 dpc embryos of zygotes pre-treated with acidic Tyrode's solution (weakened) or untreated (intact) is depicted. Arrows indicate the expected size of the band for the PCR with primers flanking the endogenous locus (WT locus) or amplifying the myc tag sequence (transgene). PCR controls from untreated embryos (WT) and without DNA template ($H_2O$) are included.

**S3 Figure**

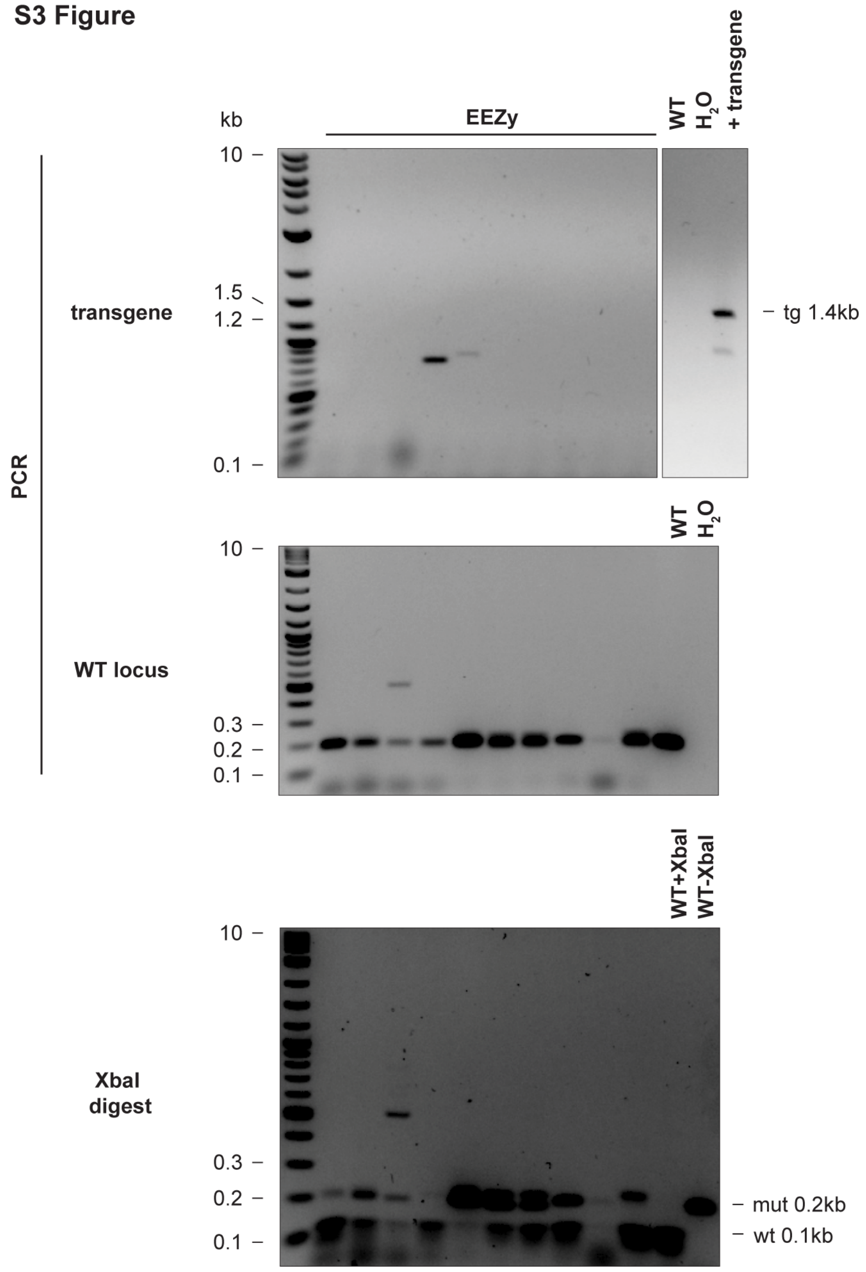


**S3 Fig. EEZy of a plasmid targeting vector attempting CRISPR-mediated transgenesis.** A circular *Gt(ROSA)26Sor* targeting vector (6 kB) harboring a Venus reporter transgene fused to a splice acceptor site (*Gt(ROSA)26Sor* SA-Venus) was electroporated into zygotes together with an Cas9 RNP targeting the *Gt(ROSA)26Sor* locus. PCR genotyping for integration of the transgene and amplification of the WT locus are depicted. PCR controls from untreated embryos (WT), without DNA template ($H_2O$) and *Gt(ROSA)26Sor SA-Venus* targeted blastocysts (+transgene) are included and the expected size for amplification of the transgene (tg) depicted. Evaluation of NHEJ at the endogenous *Gt(ROSA)26Sor* locus is shown by partial resistance to XbaI digest (mut). Controls from PCR amplicons of WT blastocysts with (WT+XbaI) and without XbaI (WT-XbaI) are included.

**S4 Figure**

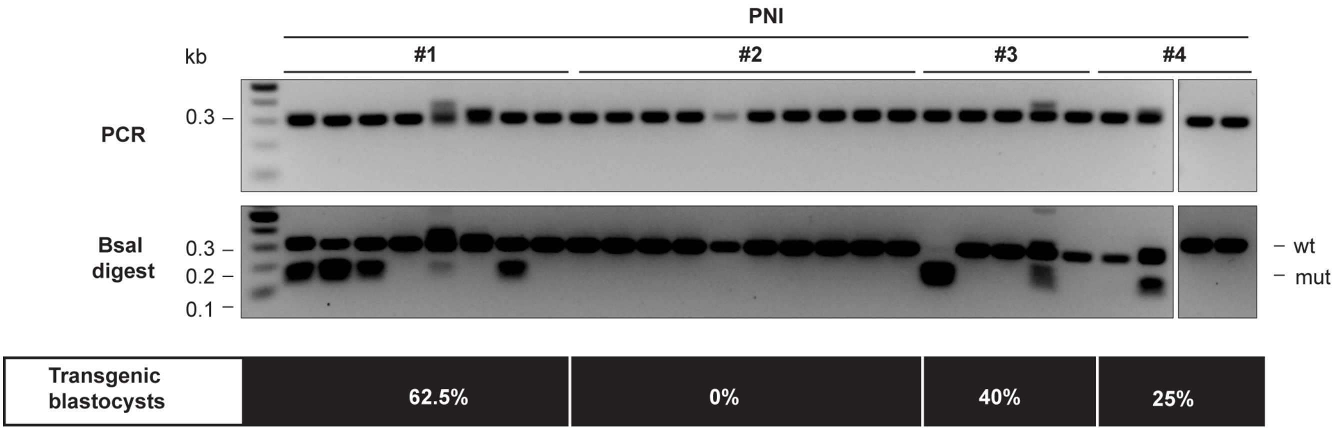


**S4 Fig. HDR efficiency in zygotes after CRISPR-PNI.** RFLP analysis of *Nphs2*-targeted blastocysts upon PNI. Results of four independent experiments (#1-4) with the percentage of transgenic blastocysts are depicted (n=4). The data are quantified in Fig 3E.

#### S1 Table. Sequences of oligonucleotides used in this study

##### S1A Table. Sequences of guide RNA and DNA repair templates

| Target | Guide RNA sequence (5'-3') | DNA repair template sequence (5'-3') |
|---|---|---|
| *Nphs2* | UUUCCAGGAGAAUUUCAGUG | CTTCTTCTAAGCAGTCTAGCCCATGTGTCCAAAGCCATCCAGTT CCTGGTGCAAACCACCATGAAGCGCCTCTTGGCACATCGGTCT CTCATTGAAATTCTCCTGGAAAGGAAGAGCATTGCCCAAGA |
| *Atp1a1* | CAUCCGAAGUGCUACAGAAG | TCTGGGAGGCAACAGCGCTACCGTAACTACACAACTCACATCA TCGTTTGGCGGTTCCTCCAGATCCTCTTCTGAGATGAGTTTTTG TTCTGTAGCACTTCGGATGCCATAAGCCAGGAAACAAAGAATG GCCCCGATCCACAGTAA |
| *Gt(ROSA) 26Sor* | ACUCCAGUCUUUCUAGAAGA | SA-Venus targeting vector |

##### S1B Table. PCR and sequencing primer and thermocycling conditions

| | Primer sequence (5'-3') | Conditions | PCR rounds |
|---|---|---|---|
| *Atp1a1* [nested] | fwd: GGTCTTAGGGCATTACGGGCT<br>rev: GACGACGACATCCTCCGCAT | 98°C 3min; 34 cycles of [95°C-20s; 59°C-20s; 72°C-30s]; 72°C-3 min | 1st |
| *Atp1a1* [WT] | fwd: CCTCCCACTACTCCCGAAT<br>rev: TCTCCTAGGTGAGTCGATTTCT | 98°C 3min; 30 cycles of [95°C-20s; 55°C-20s; 72°C-20s]; 72°C-3 min | 2nd |
| *Atp1a1* [myc] | fwd: CCTCCCACTACTCCCGAAT<br>rev: CCTCCAGATCCTCTTCTGAGAT | 98°C 3min; 30 cycles of [95°C-20s; 55°C-20s; 72°C-20s]; 72°C-3 min | 2nd |
| *Nphs2* | fwd: AGACGCTGTCTGCTACTACCGC<br>rev: TCCCCAGGCCAATGATGTCAC | 98°C 3min; 34 cycles of [95°C-20s; 67.5°C-20s; 72°C-20s]; 72°C-3 min | 2 |
| *Gt(ROSA) 26Sor* [nested] | fwd: CTCGGCTAGGTAGGGGATCG<br>rev: GCATTCCAAAAGGAACCACCTT | 98°C 3min; 38 cycles of [95°C-20s; 61.5°C-20s; 72°C-90s]; 72°C-3 min | 1st |
| *Gt(ROSA) 26Sor* [WT] | fwd: GCCTCCTGGCTTCTGAGGACCG<br>rev: TCTGTGGGAAGTCTTGTCCCTCC | 98°C 3min; 38 cycles of [95°C-20s; 66.5°C-20s; 72°C-20s]; 72°C-3 min | 2nd |
| *Gt(ROSA) 26Sor* [transgene] | fwd: GGCCTCTCGAGCCTCTAGAACTATAGTG<br>rev: CAAGCTCACAAGACCTTAGGTCAGGA | 98°C 3min; 38 cycles of [95°C-20s; 62°C-20s; 72°C-45s]; 72°C-3 min | 1 |
| *Atp1a1* | fwd: CCTCCCACTACTCCCGAAT | Sanger sequencing | N.A. |
